# Towards Greater Neuroimaging Classification Transparency via the Integration of Explainability Methods and Confidence Estimation Approaches

**DOI:** 10.1101/2022.10.06.511164

**Authors:** Charles A. Ellis, Robyn L. Miller, Vince D. Calhoun

## Abstract

The field of neuroimaging has increasingly sought to develop artificial intelligence-based models for neurological and neuropsychiatric disorder automated diagnosis and clinical decision support. However, if these models are to be implemented in a clinical setting, transparency will be vital. Two aspects of transparency are (1) confidence estimation and (2) explainability. Confidence estimation approaches indicate confidence in individual predictions. Explainability methods give insight into the importance of features to model predictions. In this study, we integrate confidence estimation and explainability approaches for the first time. We demonstrate their viability for schizophrenia diagnosis using resting state functional magnetic resonance imaging (rs-fMRI) dynamic functional network connectivity (dFNC) data. We compare two confidence estimation approaches: Monte Carlo dropout (MCD) and MC batch normalization (MCBN). We combine them with two gradient-based explainability approaches, saliency and layer-wise relevance propagation (LRP), and examine their effects upon explanations. We find that MCD often adversely affects model gradients, making it ill-suited for integration with gradient-based explainability methods. In contrast, MCBN does not affect model gradients. Additionally, we find many participant-level differences between regular explanations and the distributions of explanations for combined explainability and confidence estimation approaches. This suggests that a similar confidence estimation approach used in a clinical context with explanations only output for the regular model would likely not yield adequate explanations. We hope that our findings will provide a starting point for the integration of the two fields, provide useful guidance for future studies, and accelerate the development of transparent neuroimaging clinical decision support systems.

## Introduction

In recent years, studies have increasingly sought to develop automated diagnosis approaches using machine learning and deep learning methods for a variety of neurological and neuropsychiatric disorders like schizophrenia [1]–[3], major depressive disorder [4], [5], Alzheimer’s disease [6], [7], and others. This growth can be partially attributed to the limitations of existing clinical diagnostic approaches that are often dependent solely upon symptoms, rather than empirical biological markers, for diagnosis [8]–[10]. As many disorders can have overlapping symptoms, this is particularly problematic and can lead to delays in diagnoses and misdiagnoses. Nevertheless, if the methods being developed for automated diagnosis are ever to be implemented in a clinical setting, model transparency must be taken into consideration [11]. While there are multiple dimensions to transparency [12], there is little or no available literature on the subject of integrating two of those dimensions - model confidence estimates [13], [14] and model explainability [2][15] – into the same model for neuroimaging classification. In this study, we compare two existing approaches for estimating model confidence [13], [14]. We then examine their compatibility with two popular gradient-based explainability approaches [16], [17] within the context of resting state functional magnetic resonance imaging (rs-fMRI) classification of schizophrenia (SZ), providing guidance and a starting point for future studies seeking to develop more transparent neuroimaging models.

Within the context of automated diagnosis of neurological and neuropsychiatric disorders, and SZ in particular, a variety of modalities like magnetoencephalography (MEG) [10], electroencephalography (EEG) [8], [9], [18], magnetic resonance imaging (MRI) [19]–[21], and functional magnetic resonance imaging (fMRI) [2], [6], [22] have been used. Relative to MRI, fMRI offers insight into the links between schizophrenia and brain dynamics. Relative to modalities like MEG and EEG, fMRI is recorded at lower sampling rates and thus provides less insight into the activity of brain dynamics. However, fMRI offers significantly enhanced spatial resolution and localization relative to MEG and EEG. Additionally, fMRI has been used in many studies related to schizophrenia [23]–[26]. As such, it represents a useful modality for eventual use within the context of automated diagnosis and clinical decision support. Within the context of rs-fMRI analysis, many studies have utilized functional network connectivity (FNC) for insight into the interaction of brain networks [26]–[30], so FNC represents a useful starting point for the automated diagnosis of SZ [26].

While the use of rs-fMRI FNC for automated diagnosis of neurological disorders like SZ represents a significant opportunity within the context of clinical decision support [31], black-box automated clinical decision support systems are unlikely to be accepted by clinicians. As such, model transparency represents a critical component of the eventual implementation of clinical decision support systems. An important aspect of transparency is the capacity to provide an estimate of confidence in predictions [12]. To this end, one approach, called Monte Carlo dropout (MCD), has seen increasing use within the domain of neuroimaging classification [13]. MCD has been used in a variety of studies, including those focused on cortex parcellation [32], dynamics estimation [33], [34], and classification of autism spectrum disorder [35] and Parkinson’s disease [36]. A more recently developed alternative to MCD that has seen comparatively little use in the domain of neuroimaging classification is Monte Carlo batch normalization (MCBN) [14]. A comparison of the two methods for the domain of neuroimaging classification could provide a useful point of reference for future studies.

The use of methods like MCD and MCBN that can provide estimates of prediction confidence will greatly assist the development of transparent clinical decision support systems. However, in and of themselves, they are insufficient to the task. It is also critical that automated neuroimaging-based clinical decision support systems be explainable [11], [37]–[39]. If clinicians are to use clinical decision support systems, they are ethically obligated to be able to explain the recommendations of such systems to their patients [11]. Both explainability methods [2], [40], [41] and more recently developed interpretable models [42]–[44] have been used extensively within the domain of neuroimaging analysis. Nevertheless, with the exception of our preliminary work on this topic [1], explainability methods and approaches for estimating model confidence have, to our knowledge, not yet been integrated.

As both approaches are necessary for the long-term development of clinical decision support systems, the lack of integration of confidence estimation approaches and explainability methods remains a key gap in the current capabilities of the field. In this study, we compare MCD and the more recent MCBN approaches for estimating classifier confidence to better understand their relative utility for the domain of neuroimaging classification. We then, for the first time, integrate two popular gradient-based explainability methods with MCD and MCBN, evaluating the effects of MCD and MCBN upon the explanations and seeking to determine the best approach for integrating the two transparency approaches. It is our hope that this study will provide guidance and a helpful starting point to future studies seeking to develop more transparent neuroimaging models and clinical decision support systems.

## Methods

### Data Collection

We used the Functional Imaging Biomedical Informatics Research Network (FBIRN) dataset consisting of rs-fMRI data from 151 individuals with SZ and 160 healthy controls (HCs). The dataset has been in a number of previous studies [25], [26]. The data was collected from the University of California at Irvine, the University of California at Los Angeles, the University of North Carolina at Chapel Hill, the University of New Mexico, the University of Iowa, and the University of Minnesota. Data collection was approved by the Internal Review Boards of each study site. Six sites used 3T Siemens scanners, and 1 site used a 3T General Electric scanner for collection. T2*-weighted functional images were collected with an AC-PC aligned echo-planar imaging (EPI) sequence (TR=2s, TE=30ms, flip angle=77°, voxel size = 3.4×3.4×4mm^3^, slice gap=1mm, 162 frames, 5:24 minutes). Recordings were performed while the participants’ eyes were closed.

### Data Preprocessing

After collecting the data, we used statistical parametric mapping for preprocessing and used rigid body motion correction to account for head movement. We spatially normalized the recordings to an EPI template in the standard Montreal Neurological Institute (MNI) space, resampling to 3×3×3mm^3^. Lastly, we used a Gaussian kernel with a full width at half maximum of 6mm to smooth the recordings.

After completing general preprocessing steps, we began the feature extraction process. We applied the Neuromark automated independent component analysis (ICA) pipeline [45] of the GIFT toolbox (http://trendscenter.org/software/gift) using the Neuromark_fMRI_1.0 template (also available at http://trendscenter.org/data) to extract 53 components (ICs) associated with different brain regions and structures. The Neuromark pipeline has been used extensively across a variety of studies within the field [26], [28], [29]. We then assigned the components to 7 networks – the cerebellar (CBN), default mode (DMN), cognitive control (CCN), visual (VSN), sensorimotor (SMN), auditory (ADN), and subcortical (SCN). After extracting the ICs, we extracted the dFNC values using a sliding window approach. We used a tapered window created by convolving a rectangle of window size = 40s with a Gaussian (σ=3). We calculated Pearson’s correlation between each of the 53 ICs at each time step, resulting in 1378 dFNC features and 124 time steps per study participant. It should be noted that each of the samples can be assigned to 1 of 28 domain pairs (e.g., connectivity between CCN and VSN is CCN/VSN). After extracting each of the dFNC features, we performed feature-wise z-scoring across all subjects.

### Model Development

As shown in Figure 1, we developed a 1D-CNN architecture. We applied a 10-fold stratified shuffle split cross-validation approach from Scikit-learn with an approximately 80-10-10 training-validation-test split. Within each fold, we applied data augmentation to triple the number of training samples. This involved adding Gaussian noise (μ = 0, σ = 0.7) to two copies of each training sample. We used an Adam optimizer with an adaptive learning rate [46]. The learning rate started at 0.001 and decreased by 50% after each 15 epochs without an increase in validation accuracy (ACC). We used Kaiming He normal initialization. To address class imbalances, we used a class-weighted categorical cross-entropy. We trained the model for 100 epochs, shuffling after each epoch and using a batch size of 50. We also used a model checkpoint approach to select the model from the epoch with the highest validation accuracy. When assessing test performance, we calculated the sensitivity (SENS), specificity (SPEC), and ACC for each fold. These metrics are shown in equations 1, 2, and 3, respectively.

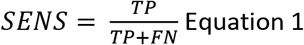

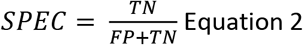

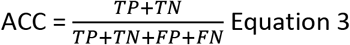

**Figure 1.**
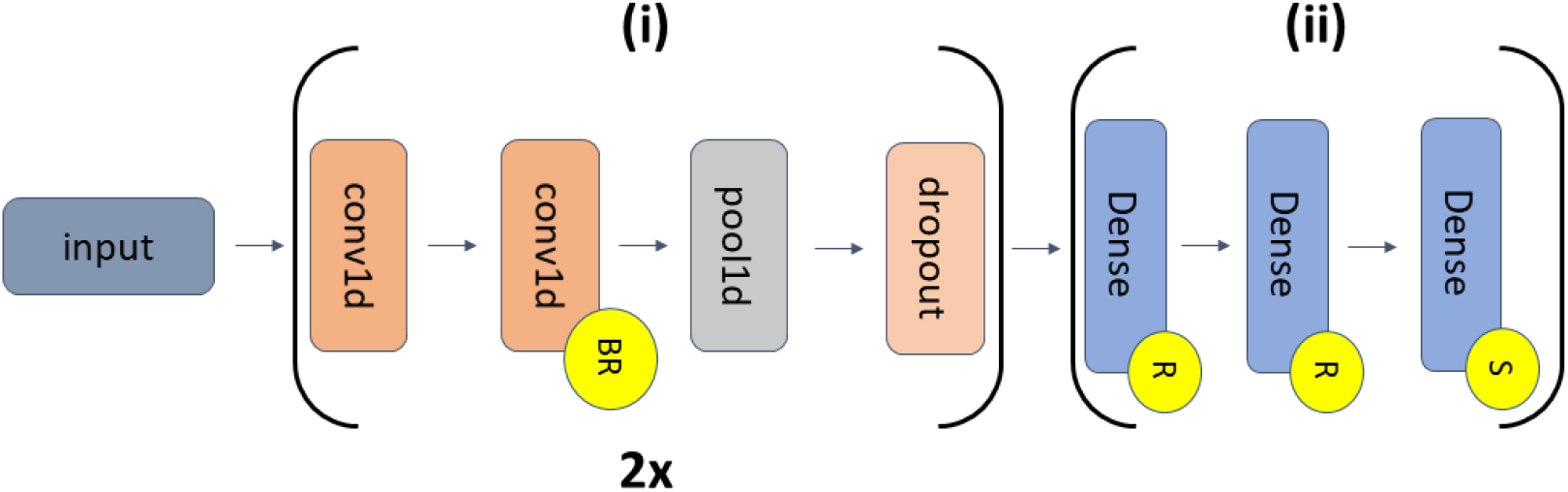
CNN Architecture. The model has sections (i) and (ii) for feature extraction and classification, respectively. Layers in section (i) are repeated twice for different hyperparameters. The first and second pairs of convolutional layers (conv1d) have a kernel size of 10 and 16 and 24 filters, respectively. Each pair of conv1d layers is followed by a max pooling layer with a pool size of 2 and spatial dropout (rates = 0.3 and 0.4). Layers in section (ii) include 3 dense layers with 10, 6, and 2 nodes. Yellow circles with “BR”, “R”, and “S” correspond to layers with batch normalization/ReLU, ReLU, and softmax activations, respectively.

Where true positives are abbreviated, “TP”, false negatives are abbreviated, “FN”, true negatives are abbreviated, “TN”, and false positives are abbreviated, “FP”.

### Monte Carlo Dropout

Monte Carlo Dropout (MCD) was first presented in [13]. At a high level, it involves the repeated reinitialization of dropout layers during testing to form a distribution of models, and thus a distribution of test predictions. It hinges upon the realization that dropout can be used to form a Bayesian approximation. In our implementation, we used 1,000 iterations of dropout during testing to form a distribution of predictions for each sample.

### Monte Carlo Batch Normalization

Similar to MCD, Monte Carlo Batch Normalization (MCBN) hinges upon the realization that another component of neural networks (i.e., batch normalizations layers) can be used to generate a Gaussian distribution of predictions. MCBN was first presented in [14]. MCBN involves several steps: (1) randomly selecting a minibatch of training data, (2) updating the model batch normalization layers based upon the minibatch of training data, (3) using the updated model to predict class probabilities for the test data, and (4) repeating steps 1 through 3 a number of times to form a distribution of predictions for each sample. We repeated MCBN for 1,000 iterations.

### Explainability

In this study, we applied two gradient-based explainability approaches: saliency and layer-wise relevance propagation.

#### Saliency

Saliency was one of the first gradient-based explainability methods [17]. It is a fairly straightforward approach that involves taking the gradient of the predicted probability of a sample belonging to a particular class with respect to each of the input features. It indicates the effect that a small change in an input feature has upon the output probability of belonging to a particular class. Larger sensitivity values correspond to a greater level of importance. Saliency has been applied in both neuroimaging studies [47] and studies involving other healthcare data types [48][49].

We applied saliency to both the original network and to the network following each iteration of MCBN and MCD, generating a distribution of values for each feature. Afterwards, we normalized the absolute value of the saliency for each study participant to make sure that the saliency summed to 100.

#### Layer-wise Relevance Propagation

Layer-wise relevance propagation (LRP) [50] is a popular gradient-based feature attribution explainability method [51]. It has been shown to produce less noisy explanations than saliency [52]. It was first developed within the context of image classification. However, because it is widely applicable to a number of deep learning architectures, it has since been applied to a number of data types, including various neuroimaging modalities [40], [53] and other healthcare data involving both time-series [54], [55] and extracted features [56].

LRP involves a series of steps. (1) A sample is passed through a network and assigned to a particular class. (2) A total relevance value of 1 is assigned to the output node corresponding to the class of the sample. (3) The relevance is propagated back through the network from layer to layer using a relevance rule until the relevance is distributed across the input space. It should be noted that depending upon the relevance rule, there can be both positive and negative relevance. Positive relevance indicates that a particular input feature provides evidence for the sample being assigned to the class to which it was assigned. Negative relevance indicates that a particular input feature gives evidence for a sample being assigned to a class other than that to which it was assigned. In this study, we used the αβ-rule to propagate only positive relevance (α = 1, β = 0). The αβ-rule is shown in equation 4.

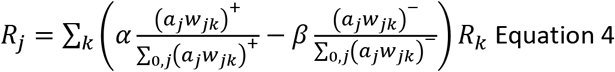

Where the relevance is split into a positive component with a coefficient α and into a negative component with a coefficient β. The variables *k* and *j* indicate a node in a deeper and shallower layer, respectively. The variables *a* and w indicate the activation associated with a particular node and the weight connecting two nodes in different layers, respectively.

We applied LRP to both the original network and to the network following each iteration of MCBN and MCD, generating a distribution of relevance values for each feature. After propagating relevance through the network in our study, we normalized the absolute relevance for each study participant to make sure that the relevance summed to 100.

### Statistical Analyses

We performed four sets of statistical analyses. The first pair of analyses were performed to gain insight into the explanations spatially, and the second pair of analyses sought insight into the temporal distribution of importance.

#### Spatial Analyses

We first sought to understand whether there were differences in the importance of individual network pairs between HCs and SZs and then sought to understand whether there were differences in the spatial distributions of importance between the basic model and the model with MCBN and MCD. (1) For insight into the spatial effects of MCBN and MCD upon the explanations for HCs relative to SZs, we summed the total importance (i.e., normalized relevance for LRP and normalized saliency for saliency) of each dFNC feature across time for each participant. We then averaged the importance within each network domain pair and across participants on a per-fold basis. We then used paired t-tests to determine whether there were differences in spatial importance distributions for HCs versus SZs across folds. We then applied FDR correction (p < 0.05). (2) For insight into the how representative the importance values associated with the regular model were of the importance distributions associated with the models with MCBN and MCD, we again summed the total importance of each dFNC feature across time for each subject. We then performed a one-sample t-test for each participant and calculated the percentage of participants for which there was a significant difference between the importance values associated with the regular model and the importance distributions of the model with MCBN and MCD. We then applied FDR correction (p < 0.05) to the values for each subject on a per-fold and per-feature basis and calculated the mean percentage of participants across folds with significant differences. We implemented these analyses for both LRP and saliency.

#### Temporal Analyses

In our temporal analyses, we first sought to understand whether there were differences in the temporal distribution of importance over time between SZs and HCs and next sought to understand whether there were differences in the temporal distribution of importance between the basic model and model with MCBN and MCD. To this end, we adapted a method presented in [15]. We used Earth mover’s distance (EMD), a distance measure comparing two densities, to calculate the distance for each participant between the importance of each dFNC feature over time with the average importance across time. A smaller EMD value indicates that the importance values are more evenly distributed across time, and a larger EMD value indicates that the importance values are more concentrated within smaller time windows. In our temporal analyses, we summed the total absolute importance for each feature on a per-participant basis and normalized that value such that the total importance summed to 100. We then compared that distribution to a distribution in which the total importance was distributed evenly over time. (1) We first sought to determine whether there were significant differences in the temporal distribution of importance across classes. We calculated the mean EMD values in each network domain pair across HCs and across SZs on a per-fold basis fold and then performed a paired t-test comparing the values for HCs and SZs. (2) We next sought to determine whether the MCD and MCBN importance distributions over time were significantly different from those of the regular model. To do this, we performed one-sample t-tests comparing the MCD and MCBN importance distributions for each dFNC feature with the importance value for the regular model. We next applied FDR correction (p < 0.05) to the values for each participant on a per-fold and per-feature basis and calculated the mean percent of participants with significant differences across folds. We repeated these analyses for both LRP and saliency.

### Analysis of Not-a-Number (NaN) Counts

While our spatial and temporal analyses did provide insight into the distribution of importance values. They overlooked an important aspect of integrating confidence estimate approaches with gradient-based explainability methods. We thought it possible that changes in model gradients associated with MCD and MCBN might adversely affect the capacity of gradients to be calculated for saliency or for relevance to be propagated for LRP. To this end, we performed two analyses. (1) We calculated the percentage of samples for each fold that returned Not-a-Number (NaN) importance values for the regular model and for at least one of the 100 iterations of MCD and MCBN. (2) We also calculated the percentage of MCD and MCBN iterations for each sample that produced NaN values across folds. In the case of LRP, relevance values might ordinarily be returned in a minority of cases if the total positive and negative relevance for a particular layer summed to zero, while saliency should not ordinarily return NaN values. The presence of a large percentage of NaN values for MCD and MCBN values would indicate that the methods adversely affected model gradients and are in some cases incompatible with gradient-based explainability methods.

## Results

In this section, we describe the results of our examination of the effects of MCD and MCBN upon model predictions and performance.

### Model Performance

Table 1 shows the mean and standard deviation of the performance of our architecture across 10 folds without a confidence estimation (i.e., “Regular”) and with MCD and MCBN. All metrics had mean levels of performance across folds that were much higher than chance-level, with average accuracies around 75%. Interestingly, the model sensitivity and specificity were most balanced for the regular model. In contrast, MCBN and MCD seemed to favor either sensitivity or specificity. MCBN favored SPEC much more than it favored SENS, while MCD favored SENS much more than it favored SPEC. Nevertheless, mean ACC for the model with MCBN was similar to that of the regular model, and the mean ACC was slightly lower for the model with MCD.

**Table 1.** Model Performance Results.

| Table 1. Model Performance Results. |  |  |  |
| --- | --- | --- | --- |
|  | SENS | SPEC | ACC |
| Regular | 75.63 ± 14.64 | 74.38 ± 11.34 | 75.00 ± 07.26 |
| MCBN | 70.63 ± 15.32 | 79.35 ± 06.87 | 75.00 ± 09.06 |
| MCD | 78.13 ± 13.76 | 68.13 ± 13.76 | 73.13 ± 07.93 |

### Distribution of Predictions

Figure 2 shows the distribution of sample predictions across folds for the regular model without confidence estimation and for the model with MCD and MCBN. The classifier generally predicted either very high or very low probabilities, even in some cases of misclassification. However, particularly among HCs, there were some samples that fell along a gradient of predictions closer to the 50% line. The range of MCD sample predictions was much wider than the range of MCBN sample predictions. With the exception of samples that were already near the decision boundary, the standard deviation of the predictions for MCBN was typically very small. Lastly, in a large number of instances, MCD seemed to move the sample predictions closer to the 50% decision boundary.

**Figure 2.**
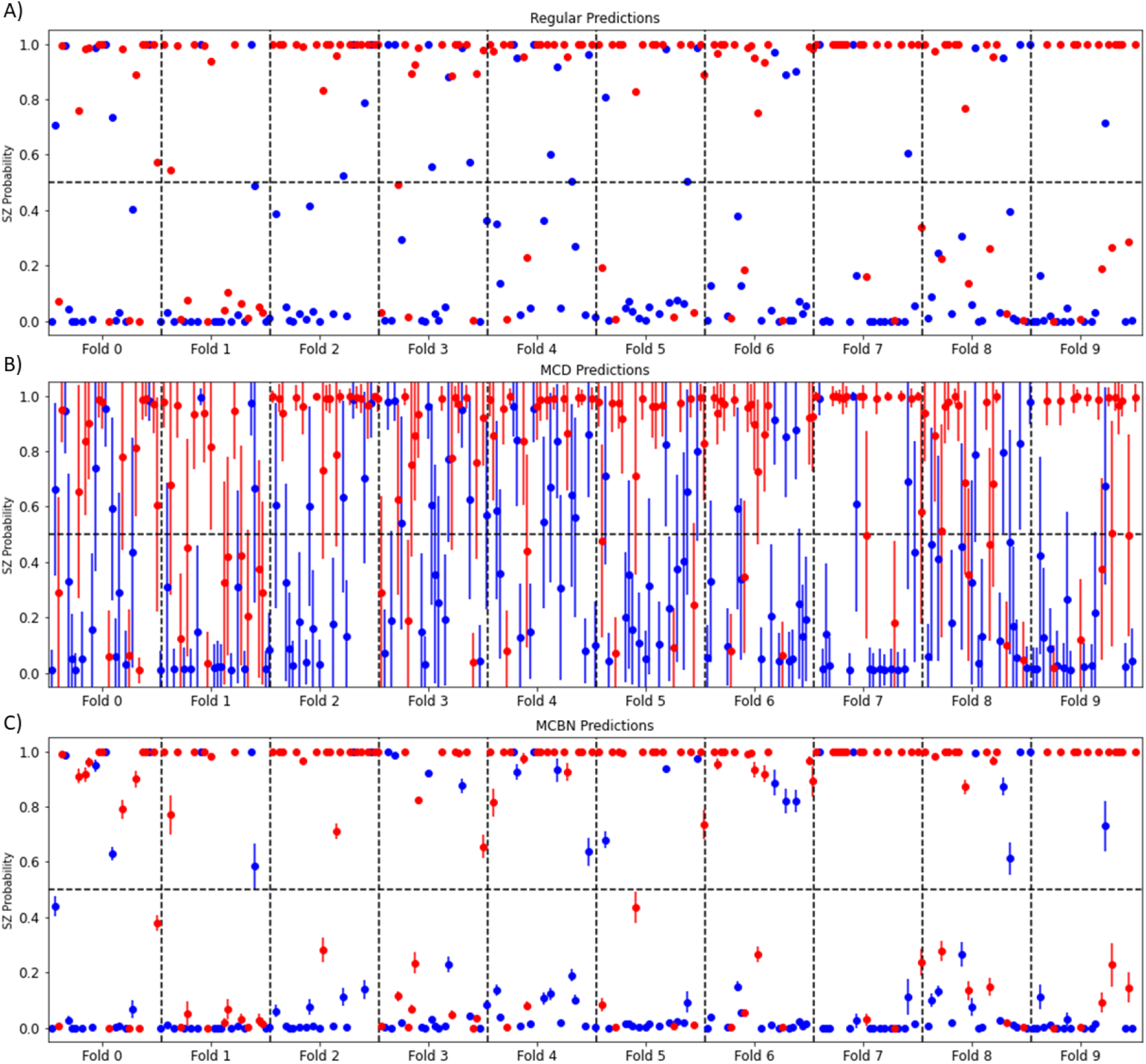
Distributions of Sample Predictions. Panel A shows the model predictions without confidence estimation. Panels B) and C) show the model predictions with MCD and MCBN, respectively. The predictions for samples with true labels of HC and SZ are shown in blue and red respectively, and samples are aligned in the same order across panels such that each panel can be visually compared. It should be noted that the points in Panels B and C reflect the mean of predictions, and the error lines reflect one standard deviation above and below the mean. Samples are grouped from left to right based on their folds, with a black dashed vertical line separating samples for each fold. The y-axis reflects the probability of a sample belonging to the SZ class, and the black dashed horizontal line indicates the 50% boundary point between classes.

### Spatial Analysis

Figures 3 and 4 show the spatial distributions of importance for LRP and saliency, respectively (i.e., the sum of the absolute value of the importance across all time steps). For both LRP and saliency, there were several brain network pairs with high levels of importance across nodes for the classification, including CBN/SCN and SMN/SCN. Other network pairs, including VSN/SMN, VSN/SCN, and VSN, had high levels of importance for specific brain regions that they contained. Specifically, (1) the interactions of the VSN with the postcentral gyrus of the SMN, (2) the interactions of the VSN with the thalamus of the SCN, and (3) the interactions of the other VSN brain regions with the cuneus region were important. For LRP, the interactions of the SMN with the inferior parietal lobule of the CCN were important. While saliency did not identify this particular interaction as being as important as LRP, it indicated that the interaction of two SMN brain regions with the inferior parietal lobule of the CCN was importance.

**Figure 3.**
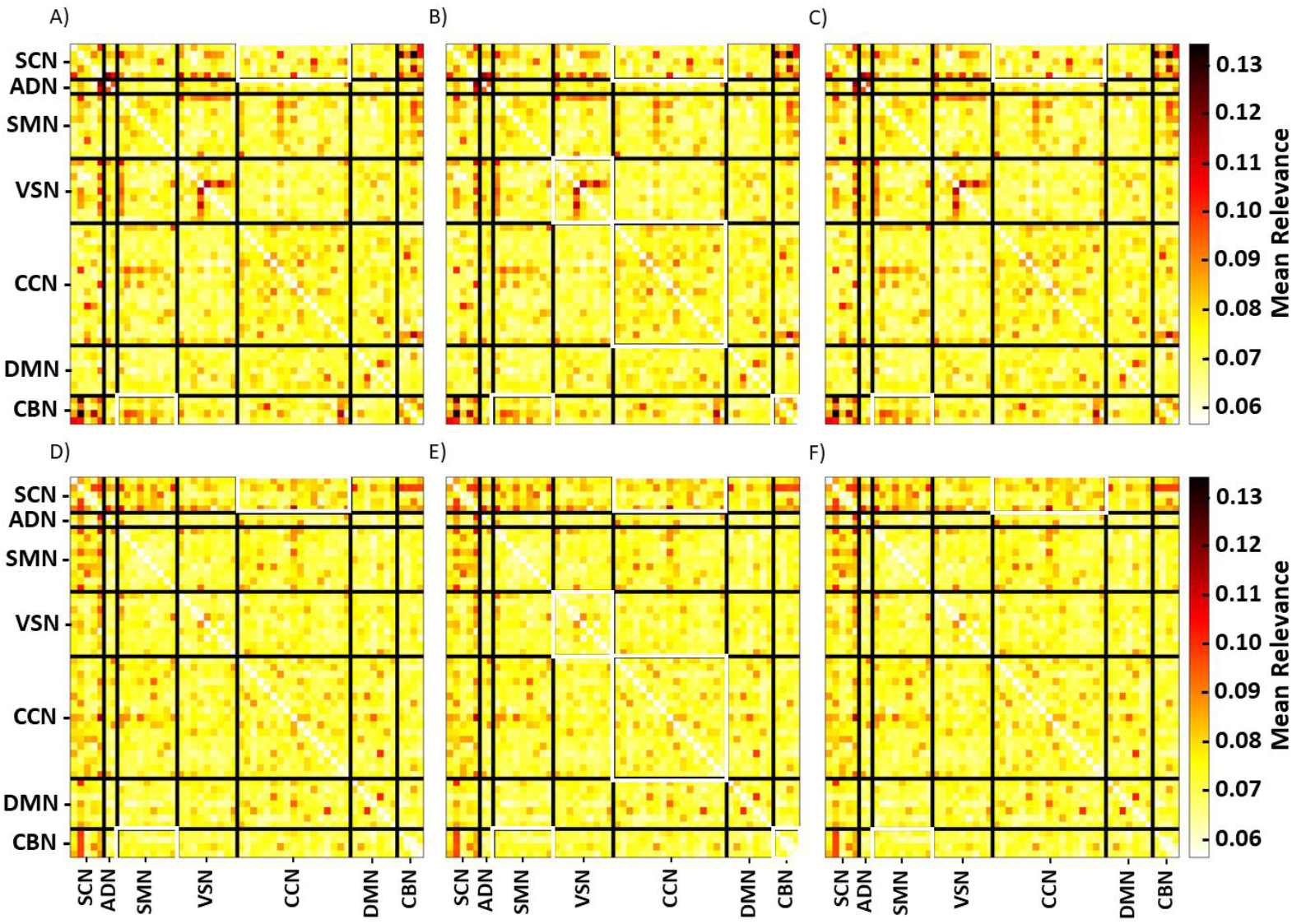
Mean of Total LRP Relevance Across All Timesteps. Panels A, B, and C reflect the mean relevance of the regular model, the model with MCD, and the model with MCBN for HCs. Panels D, E, and F show the same values for SZs. Networks are included on the x- and y-axes and are separated by black lines. Network pairs surrounded by white boxes have statistically significant differences between HCs and SZs. Lastly, all panels share the color bars to the right of Panels C and F.

**Figure 4.**
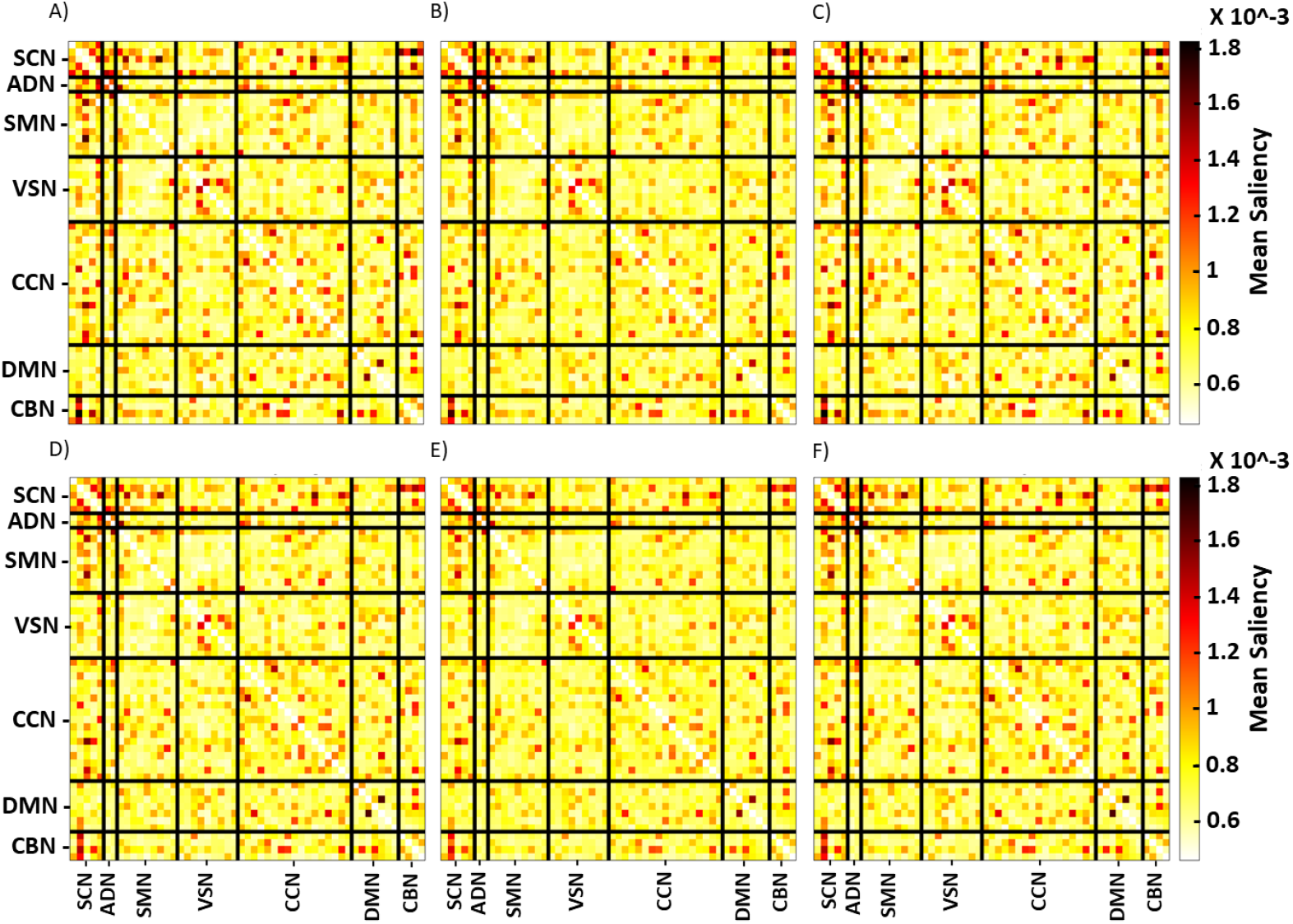
Mean of Total Saliency Across All Timesteps. Panels A, B, and C reflect the mean saliency of the regular model, the model with MCD, and the model with MCBN for HCs. Panels D, E, and F show the same values for SZs. Networks are included on the x- and y-axes and are separated by black lines. All panels share the color bars to the right of Panels C and F.

Additionally, while there were no significant class-specific differences in saliency values, there were some visible differences in LRP relevance between HCs and SZs. These visible differences were present in the SCN/CBN, the SCN/SCN, the SCN/SMN, the VSN/SMN, the SMN/CBN, the CCN/CBN, and the CBN/CBN. Our statistical analysis of class-specific importance found differences in LRP relevance between HCs and SZs for CCN/SCN and CBN/SMN for both the regular model and model with MCBN. In contrast, there were more network pairs with class-specific relevance differences in models with MCD – VSN/VSN, CCN/SCN, CCN/CCN, CBN/SMN, and CBN/CBN.

For both LRP and saliency, the overall mean importance across folds seemed to be similar across the regular model, the model with MCD, and the model with MCBN. This indicates that at a high level the three methods yield similar levels of importance. However, Figure 5 provides some higher resolution insight. It shows the average percentage of samples per fold for which there were significant differences (p < 0.05) between the model with regular LRP and the model with MCBN and MCD. Importantly, for saliency, the majority of samples had regular importance values that were away from the means of the importance values with MCD and with MCBN. Importantly, there were some differences in the percentages of samples for regular versus MCD and for regular versus MCBN in a number of network pairs. Additionally, for LRP, a large number of samples in each fold had significant differences. This was particularly the case for the regular LRP values versus the MCD LRP values, and it was true to a lesser extent in the case for the regular LRP values versus the MCBN LRP values. It should be noted, however, that the use of MCD and MCBN seemed to have a larger effect upon the saliency values than upon the LRP values.

**Figure 5.**
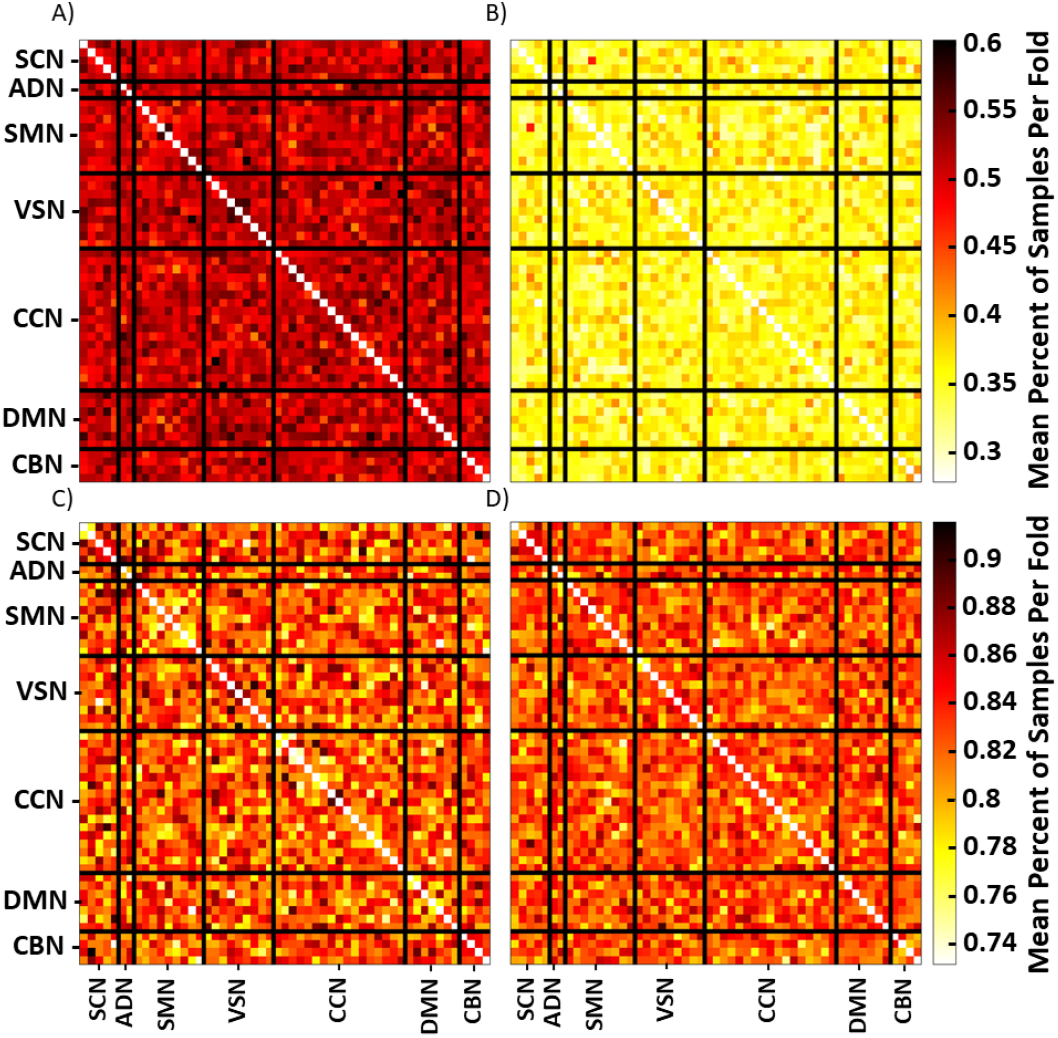
Sample Level Differences in Spatial Importance. Panels A and B show the mean percent of samples per fold with differences (p < 0.05) between their regular relevance values and their MCD and MCBN relevance distributions, respectively. Panels C and D show the mean percent of samples per fold with differences (p < 0.05) between their regular saliency values and their MCD and MCBN saliency distributions, respectively. The color bars to the right of Panels B and D are shared by Panels A and B and Panels C and D, respectively.

### Temporal Analysis

Figures 6 and 7 show the mean EMD across folds for LRP relevance and saliency, respectively. Higher EMD values indicate that importance is more concentrated temporally, while lower EMD values indicate that importance is more uniformly dispersed over time. Interestingly, for both LRP and saliency, importance is more concentrated temporally in SZs than in HCs across the regular, MCD, and MCBN implementations. This is somewhat unexpected for saliency given that the spatial analyses did not find class-specific differences in importance. Additionally, it is odd that the MCD relevance values tend to be more concentrated temporally than those for the regular and MCBN implementations, while the MCD saliency values seem to be less concentrated temporally that those of the regular and MCBN implementations. The regular and MCBN implementations generally seem to have similar values across both LRP and saliency. It should be noted that while these differences are distinct visually, they are not statistically significant. This indicates that while the differences in the mean EMD can be visualized, the distribution of EMD values do overlap to a degree.

**Figure 6.**
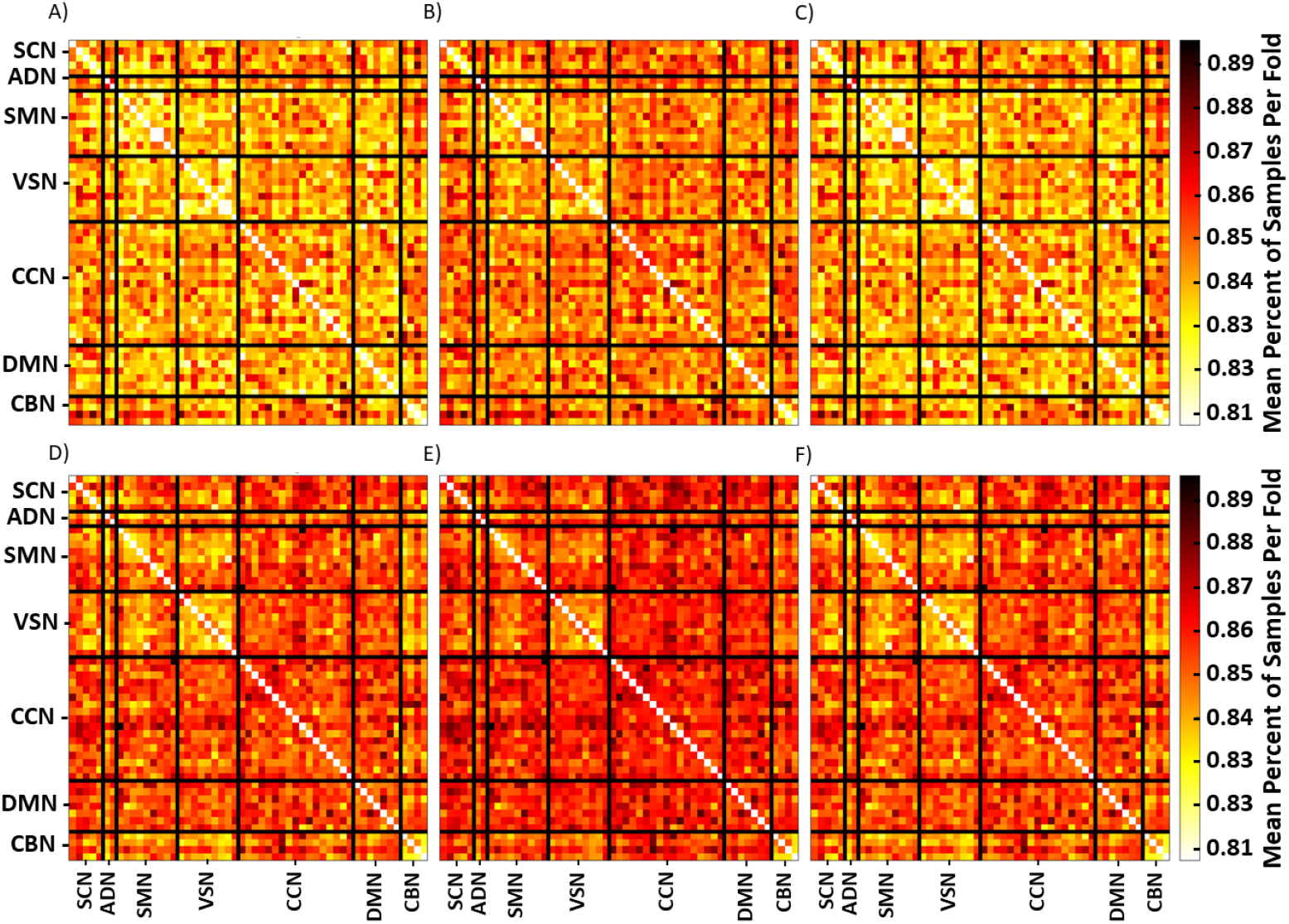
Mean of Relevance EMD over Time. Panels A, B, and C reflect the mean EMD of the regular model, the model with MCD, and the model with MCBN for HCs. Panels D, E, and F show the same values for SZs. Networks are included on the x- and y-axes and are separated by black lines. All panels share the color bars to the right of Panels C and F.

**Figure 7.**
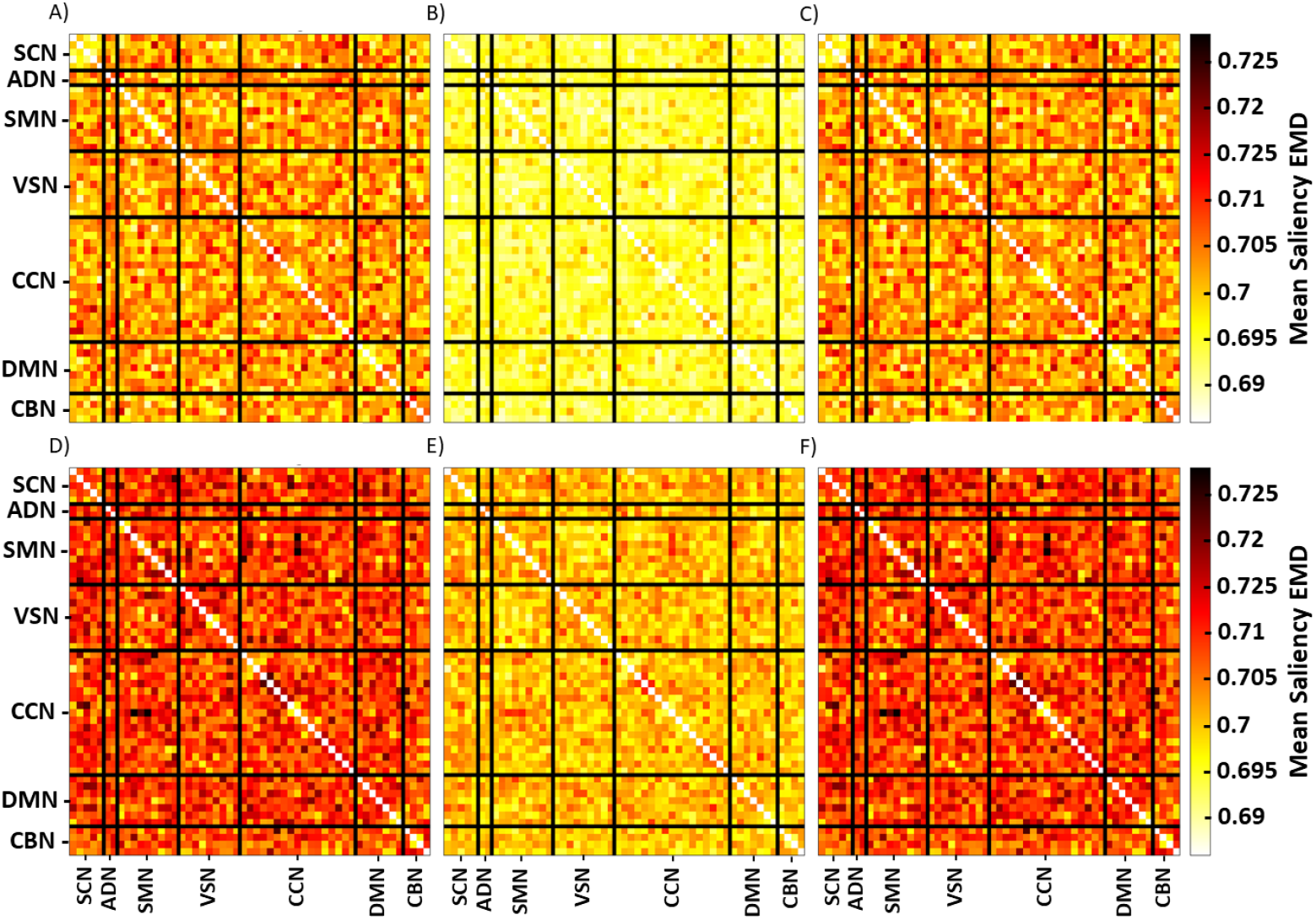
Mean of Saliency EMD over Time. Panels A, B, and C reflect the mean EMD of the regular model, the model with MCD, and the model with MCBN for HCs. Panels D, E, and F show the same values for SZs. Networks are included on the x- and y-axes and are separated by black lines. All panels share the color bars to the right of Panels C and F.

Figure 8 shows the percent of samples per fold with significant difference between the EMD of the importance for the regular model and with the MCD and MCBN models. Interestingly, for both LRP and saliency, the MCBN importance distributions were further from the importance values of the regular model than the MCD importance values. Additionally, the number of participants with differences was greater for saliency than for LRP.

**Figure 8.**
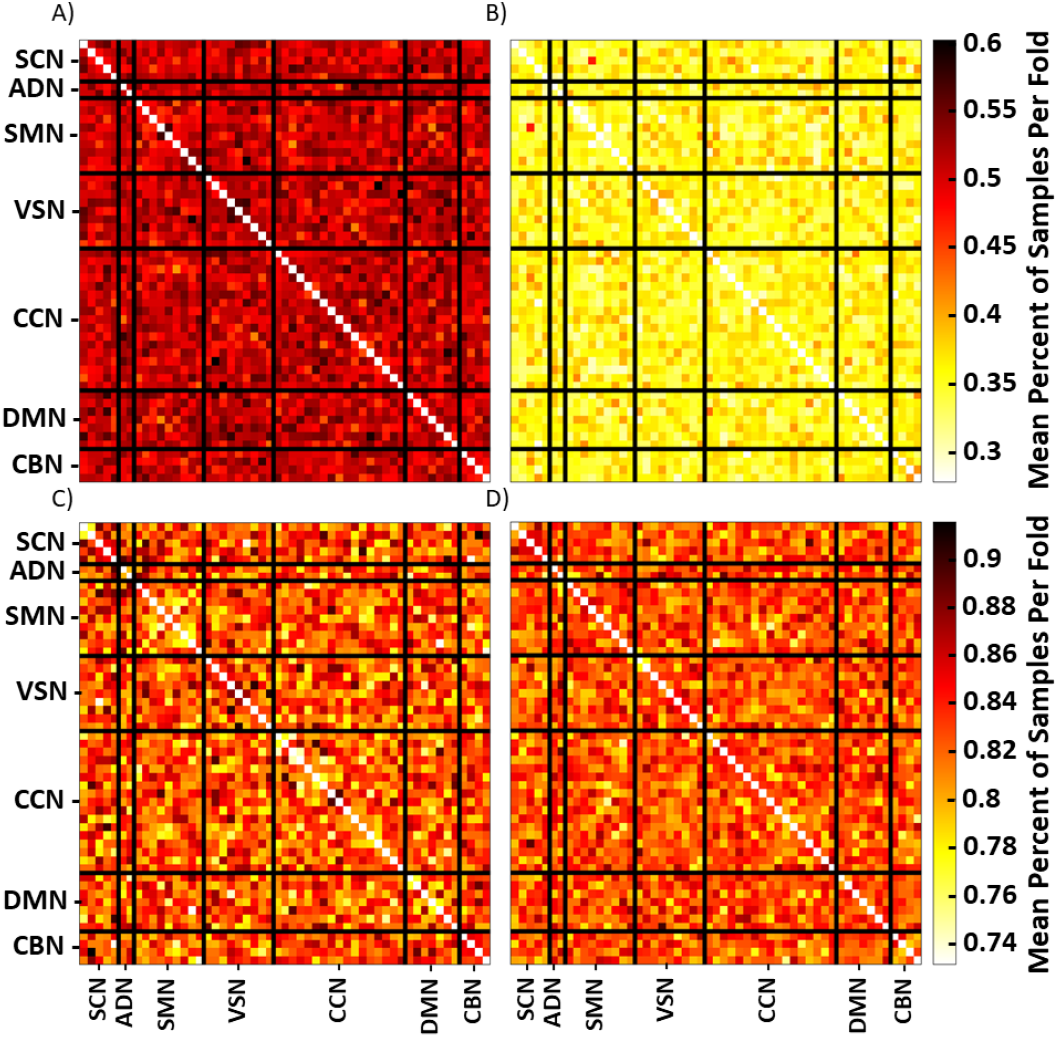
Sample Level Differences in Temporal Importance Distributions. Panels A and B show the mean percent of samples per fold with differences (p < 0.05) between their regular relevance EMD values and their MCD and MCBN EMD distributions, respectively. Panels C and D show the mean percent of samples per fold with differences (p < 0.05) between their regular saliency EMD values and their MCD and MCBN saliency EMD distributions, respectively. The color bars to the right of Panels B and D are shared by Panels A and B and Panels C and D, respectively.

### NaN Value Analysis

Lastly, it is important to consider the effects of the confidence estimation approaches upon the viability of the explainability methods. As such, we examined the number of samples and iterations per sample of MCD and MCBN that resulted in the production of NaN values. These results are shown in Figure 9. Interestingly, saliency with the regular model and with MCBN did not return any NaN values, while LRP with the regular model and with MCBN did on one occasion in two of ten folds. For a significant portion of samples, the model with MCD returned at least one iteration of NaN values for both explainability methods. The percentage of NaN values ranged from around zero to ten percent of iterations for most folds across, with some folds reaching much higher levels (e.g., up to 50% for LRP).

**Figure 9.**
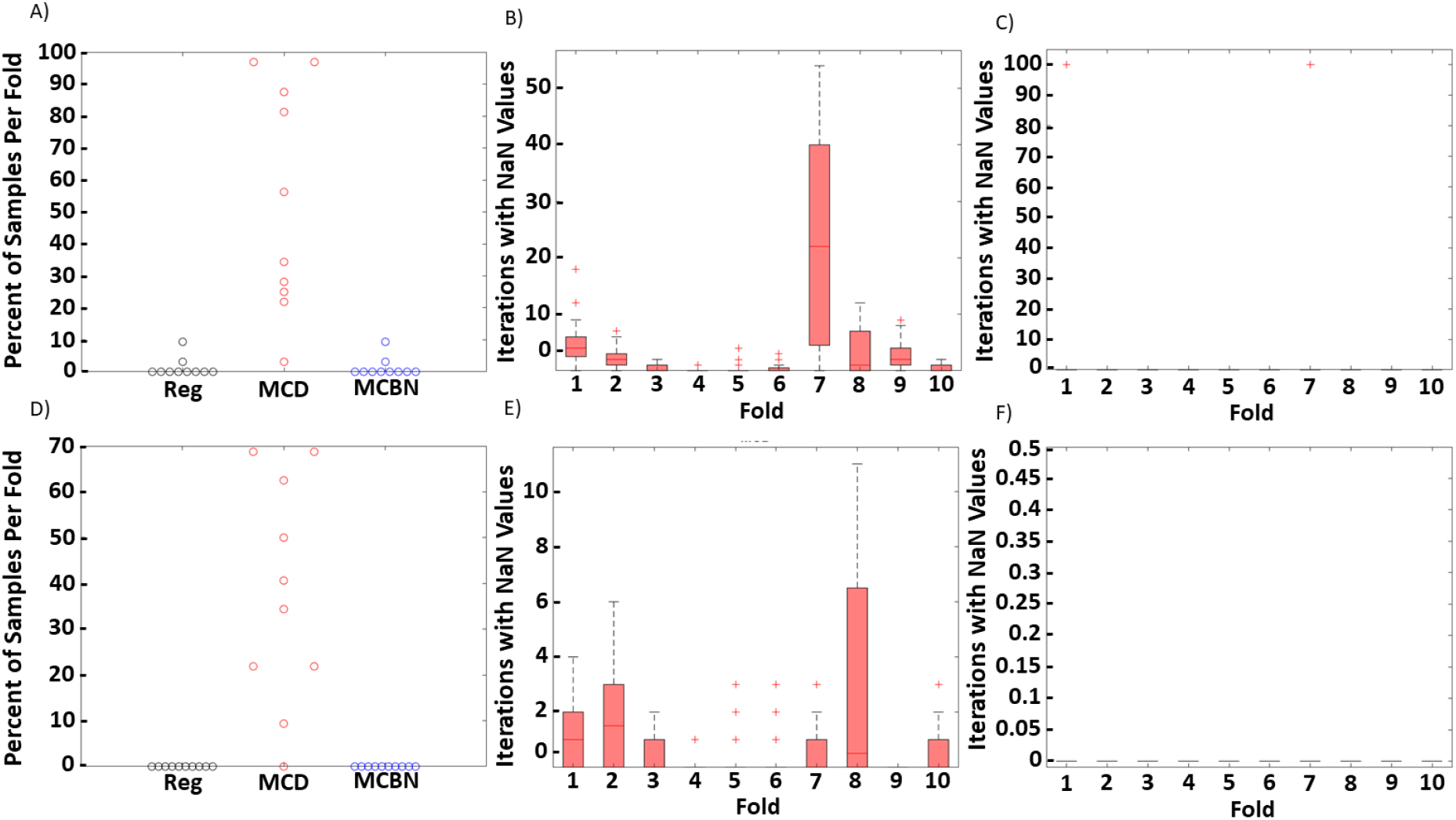
Distribution of NaN Values. Panels A and D show the percent of samples per fold with at least one NaN value for LRP and saliency, respectively. The values for the regular model, the model with MCD, and the model with MCBN are shown in black, red, and blue, respectively. Panels B and C show the percent of iterations per sample of each fold that produced NaN LRP relevance values for MCD and MCBN, respectively. Panels E and F show the percent of iterations per sample of each fold that produced NaN saliency values for MCD and MCBN, respectively.

## Discussion

In this section, we discuss the performance results of our model and the results of the effects of MCD and MCBN upon the model predictions. We then discuss the effects of the respective methods upon output explanations. Lastly, we discuss limitations and next steps related to our work.

### Model Performance

Overall model performance was well above chance-level. Additionally, while MCBN and MCD were able to provide distributions of predictions for each sample, they seemed to destabilize the SENS and SPEC of the model. This was a surprising finding relative to previous studies that have used MCD and MCBN [57], suggesting that the utility of the methods for improving classifier performance may be somewhat dependent upon the utilized model and dataset. Relative to the performance of the models developed in [2] that had accuracies ranging from 50% to 83%, our model performance was on par to slightly lower. This could partially be related to the significantly larger dataset size used in [2]. Additionally, our model performance was below that of [47], which used a novel form of extracted features.

### Distribution of Predictions

Our model generally had somewhat polarized predictions for each class, with extremely high probabilities of samples belonging to one class and extremely low probabilities of samples belonging to the other class. While this was the case, there were a number of samples that were misclassified or that were predicted to be closer to the decision boundary line. The predictions of MCBN tended to be more stable across iterations. In contrast, the predictions of MCD generally varied widely. Thus, it seems that for our data and model the use of dropout during testing more greatly affects predictions than changes in the batch normalization layer values. Considering the distributions of predictions as well as the difference in model performance, we can conclude that MCBN offered a better approach.

### Spatial Analysis

We identified a number of brain network pairs useful to the classification of SZ and HCs. Previous studies have found widespread effects of SZ upon the CBN [2][15][27], SMN [15], and SCN [2] similar to our results. Additionally, some studies have identified differences in the VSN/VSN [25][58], VSN/SCN [25][59], and VSN/SMN [25][60] as important to differentiating SZ from healthy individuals. Specifically, changes in connectivity between the VSN and thalamus of the SCN have been identified [59]. These findings support the overall reliability of our model relative to previous studies.

Our findings on the differences between class-specific LRP and saliency results were largely unsurprising. Given the characteristics of each of the methods. LRP is able to provide class-specific relevance values when using the αβ-rule, so we would expect to see differences in the relevance distribution between classes. In contrast, saliency just shows the gradient associated with the classification, and it is likely that specific regions will have similar gradients across classes.

Differences between the results comparing the regular model with MCD and with MCBN could potentially be attributed to the effects that they had upon the model shown in panels C and F of Figure 5. The regular model and model with MCBN had relatively little adverse effects upon the explanations. For LRP, although we used the αβ-rule with α = 1 and β = 0, the rule can still sometimes pass backwards both positive and negative values, and in a minority of occasions the values in the denominator of Equation 4 may have summed to a total relevance of zero. In contrast to the regular model and model with MCBN, MCD frequently resulted in the returning of NaN values for explanations. As such, although MCD can sometimes improve model performance, the use of multiple layers with dropout can, under some circumstances, effectively destroy model gradients, making it incompatible with gradient-based explainability methods. This finding could also explain the relatively high variance of model predictions that we identified for MCD, and it offers a key guidance for future studies seeking to integrate confidence estimation methods with gradient-based explainability methods. Namely, MCD is unlikely to be compatible with gradient-based explainability methods.

### Temporal Analysis

We found that generally SZs had more temporally concentrated importance values than HCs. This would indicate that the aberrant effects of SZ upon brain network interactions tend to be temporally localized. This finding is supported by [15], which found similar effects of SZ upon attention values of a long short-term memory network. Additionally, this finding is related to those of other studies that have found effects of SZ upon brain network dynamics [24], [26].

At a high level, the temporal distribution of importance for the regular model and the model with MCBN tended to be more similar to one another than the temporal distribution of the model with MCD. However, on a more fine-grained, per-participant basis, there tended to be a difference between the EMD values associated with the regular model and the EMD distributions for both the model with MCD and with MCBN. Additionally, these differences were stronger for saliency than for LRP.

### Recommendations for Integration of Confidence Estimation Approaches and Explainability Methods

While MCD has seen more widespread use than MCBN, its effects upon model gradients can prevent its effective integration with explainability methods. The number of iterations in which NaN values are produced for MCD is generally not the majority of iterations. However, it would likely be necessary to discard a number of MCD iterations in any sort of implementation setting. As such, MCBN represents the more viable of the two methods for combination with gradient-based explainability methods. Within the context of applying MCBN with explainability methods, there are not large differences between explanations with MCBN relative to explanations for a regular model when those explanations are averaged across individuals and folds. As such, during model development, it may be unhelpful to integrate the two methods. However, we also found that on a per-individual basis, neither the spatial nor the temporal distribution of importance values associated with a traditional deep learning model tend not to be representative of the overall distribution that results when the model is combined with MCBN. As such, in the context of clinical implementation in which clinicians will be examining explanations on an individual patient basis and in which they will likely be evaluating model confidence estimates, it will be more necessary to output explanations for each iteration of confidence estimation approaches if the confidence estimates being examined by clinicians are to be properly explained. Additionally, some explainability methods, like saliency relative to LRP, tend to be noisier than others [52]. This tendency seems to be amplified when combined with confidence estimation approaches, as we found that generally the distributions of MCBN saliency were significantly different from their corresponding regular model importance values in more individuals than were those of LRP.

### Limitations and Next Steps

While explainability is needed for the development of clinical decision support systems [11], [37]–[39], a number of researchers have indicated that current explainability approaches are insufficient for use in a clinical setting [38][61]. There are valid concerns associated with their critiques. Nevertheless, for the purposes of this study, we only sought to provide a starting point to the integration of confidence estimation approaches and explainability methods. Further developments will be needed within the context of both confidence estimation and explainability in future years. However, they will eventually need to be integrated, and it is better that the field begin considering that integrative process sooner rather than later. While existing confidence estimation methods have had some popularity within the context of neuroimaging classification in recent years, they require repeated predictions and can be computationally intensive. Their repeated combination when directly paired with repeated output of explanations can be doubly intensive to the point of impracticality. This is, again, an example of how novel developments will be needed for both fields in the coming years.

## Conclusion

In this study, we combined model confidence estimation approaches with explainability methods for the first time to help address the need for greater transparency in neuroimaging-based clinical decision support systems. We used two confidence estimation approaches – MCD and MCBN. We further combined the two approaches with saliency and LRP for explainability. Our findings indicate that MCBN obtains comparable or better classification performance than MCD. Additionally, we found that MCD often adversely affected model gradients, while MCBN did not. We also uncovered spatial and temporal effects of SZ upon brain activity using our approach. It is our hope that this study will provide a starting point to the field on the integration of confidence estimation and explainability methods, provide useful guidance for future studies, and accelerate the development of transparent neuroimaging clinical decision support systems.

## References

[1] C. A. Ellis, R. L. Miller, and V. D. Calhoun, “An Approach for Estimating Explanation Uncertainty in fMRI dFNC Classification,” bioRxiv, 2022.

[2] W. Yan et al., “Discriminating schizophrenia using recurrent neural network applied on time courses of multi-site FMRI data,” EBioMedicine, vol. 47, pp. 543–552, Sep. 2019, doi: 10.1016/j.ebiom.2019.08.023.

[3] B. Rashid et al., “Classification of schizophrenia and bipolar patients using static and dynamic resting-state fMRI brain connectivity,” Neuroimage, vol. 134, pp. 645–657, 2016, doi: 10.1016/j.neuroimage.2016.04.051.

[4] B. Sen, B. Mueller, B. Klimes-Dougan, K. Cullen, and K. K. Parhi, “Classification of Major Depressive Disorder from Resting-State fMRI,” Proc. Annu. Int. Conf. IEEE Eng. Med. Biol. Soc. EMBS, no. Mdd, pp. 3511–3514, 2019, doi: 10.1109/EMBC.2019.8856453.

[5] J. Y. Chun, M. S. E. Sendi, J. Sui, D. Zhi, and V. D. Calhoun, “Visualizing Functional Network Connectivity Difference between Healthy Control and Major Depressive Disorder Using an Explainable Machine-learning Method,” in 2020 42nd Annual International Conference of the IEEE Engineering in Medicine &amp; Biology Society (EMBC), 2020, pp. 955–960, doi: 10.1109/BIBE50027.2020.00162.

[6] E. Challis, P. Hurley, L. Serra, M. Bozzali, S. Oliver, and M. Cercignani, “Gaussian process classification of Alzheimer’s disease and mild cognitive impairment from resting-state fMRI,” Neuroimage, 2015, doi: 10.1016/j.neuroimage.2015.02.037.

[7] S. Liu, S. Liu, W. Cai, S. Pujol, R. Kikinis, and D. Feng, “EARLY DIAGNOSIS OF ALZHEIMER’S DISEASE WITH DEEP LEARNING,” 2014.

[8] C. A. Ellis, A. Sattiraju, R. L. Miller, and V. D. Calhoun, “Examining Effects of Schizophrenia on EEG with Explainable Deep Learning Models,” in bioRxiv, 2022, pp. 5–8.

[9] C. A. Ellis, A. Sattiraju, R. Miller, and V. Calhoun, “Examining Effects of Schizophrenia on EEG with Explainable Deep Learning Models,” bioRxiv, pp. 5–8, 2022.

[10] T. J. Gawne et al., “A multimodal magnetoencephalography 7 T fMRI and 7 T proton MR spectroscopy study in first episode psychosis,” npj Schizophr., vol. 6, no. 1, pp. 1–9, 2020, doi: 10.1038/s41537-020-00113-4.

[11] J. Amann, A. Blasimme, E. Vayena, D. Frey, and V. I. Madai, “Explainability for artificial intelligence in healthcare: a multidisciplinary perspective,” BMC Med. Inform. Decis. Mak., vol. 20, no. 1, pp. 1–9, 2020, doi: 10.1186/s12911-020-01332-6.

[12] U. Bhatt et al., “Uncertainty as a Form of Transparency: Measuring, Communicating, and Using Uncertainty,” AIES 2021 - Proc. 2021 AAAI/ACM Conf. AI, Ethics, Soc., pp. 401–413, 2021, doi: 10.1145/3461702.3462571.

[13] Y. Gal and Z. Ghahramani, “Dropout as a Bayesian approximation: Representing model uncertainty in deep learning,” 33rd Int. Conf. Mach. Learn. ICML 2016, vol. 3, pp. 1651–1660, 2016.

[14] M. Teye, H. Azizpour, and K. Smith, “Bayesian uncertainty estimation for batch normalized deep networks,” in 35th International Conference on Machine Learning, ICML 2018, 2018, vol. 11, pp. 7824–7833.

[15] M. Rahman et al., “Interpreting models interpreting brain dynamics,” Sci. Rep., 2022.

[16] S. Bach, A. Binder, G. Montavon, F. Klauschen, K. R. Müller, and W. Samek, “On pixel-wise explanations for non-linear classifier decisions by layer-wise relevance propagation,” PLoS One,vol. 10, no. 7, Jul. 2015, doi: 10.1371/journal.pone.0130140.

[17] K. Simonyan, A. Vedaldi, and A. Zisserman, “Deep Inside Convolutional Networks: Visualising Image Classification Models and Saliency Maps,” Dec. 2013, [Online]. Available: http://arxiv.org/abs/1312.6034.

[18] L. Zhang, “EEG Signals Classification Using Machine Learning for the Identification and Diagnosis of Schizophrenia,” Proc. Annu. Int. Conf. IEEE Eng. Med. Biol. Soc. EMBS, pp. 4521–4524, 2019, doi: 10.1109/EMBC.2019.8857946.

[19] A. V. Lebedev et al., “Random Forest ensembles for detection and prediction of Alzheimer’s disease with a good between-cohort robustness,” NeuroImage Clin., 2014, doi: 10.1016/j.nicl.2014.08.023.

[20] M. Böhle, F. Eitel, M. Weygandt, and K. Ritter, “Layer-Wise Relevance Propagation for Explaining Deep Neural Network Decisions in MRI-Based Alzheimer’s Disease Classification,” Front. Aging Neurosci., vol. 11, Jul. 2019, doi: 10.3389/fnagi.2019.00194.

[21] D. Wood, J. Cole, and T. Booth, “NEURO-DRAM: a 3D recurrent visual attention model for interpretable neuroimaging classification,” Oct. 2019, [Online]. Available: http://arxiv.org/abs/1910.04721.

[22] M. S. E. Sendi, J. Y. Chun, and V. D. Calhoun, “Visualizing functional network connectivity difference between middle adult and older subjects using an explainable machine-learning method,” in Proceedings - IEEE 20th International Conference on Bioinformatics and Bioengineering, BIBE 2020, 2020, pp. 955–960, doi: 10.1109/BIBE50027.2020.00162.

[23] J. Zheng, X. Wei, J. Wang, H. Lin, H. Pan, and Y. Shi, “Diagnosis of Schizophrenia Based on Deep Learning Using fMRI,” Comput. Math. Methods Med., 2021, doi: 10.1155/2021/8437260.

[24] C. A. Ellis, M. S. E. Sendi, R. L. Miller, and V. D. Calhoun, “An Unsupervised Feature Learning Approach for Elucidating Hidden Dynamics in rs-fMRI Functional Network Connectivity,” in 2022 44th Annual International Conference of the IEEE Engineering in Medicine & Biology Society (EMBC), 2022, pp. 4449–4452.

[25] C. A. Ellis, M. S. E. Sendi, E. P. T. Geenjaar, S. M. Plis, R. L. Miller, and V. D. Calhoun, “Algorithm-Agnostic Explainability for Unsupervised Clustering,” pp. 1–22, 2021, [Online]. Available: http://arxiv.org/abs/2105.08053.

[26] M. S. E. Sendi et al., “Aberrant Dynamic Functional Connectivity of Default Mode Network in Schizophrenia and Links to Symptom Severity,” Front. Neural Circuits, vol. 15, no. March, pp. 1–14, 2021, doi: 10.3389/fncir.2021.649417.

[27] M. Liang et al., “Widespread functional disconnectivity in schizophrenia with resting-state functional magnetic resonance imaging,” Neuroreport, vol. 17, no. 2, pp. 209–213, 2006, doi: 10.1097/01.wnr.0000198434.06518.b8.

[28] M. S. E. Sendi, C. A. Ellis, R. L. Milller, D. H. Salat, and V. D. Calhoun, “The relationship between dynamic functional network connectivity and spatial orientation in healthy young adults,” bioRxiv, 2021.

[29] M. S. E. Sendi et al., “The link between brain functional network connectivity and genetic risk of Alzheimer’s disease,” bioRxiv, 2021, doi: 10.1002/alz.050101.

[30] C. A. Ellis, M. L. Sancho, R. Miller, and V. Calhoun, “Exploring Relationships between Functional Network Connectivity and Cognition with an Explainable Clustering Approach,” in bioRxiv, 2022, pp. 23–26.

[31] M. Pedersen, K. Verspoor, and M. Jenkinson, “Artificial intelligence for clinical decision support in neurology,” Brain Commun., pp. 1–11, 2020, doi: 10.1093/braincomms/fcaa096.

[32] L. Z. J. Williams, A. Fawaz, M. F. Glasser, A. D. Edwards, and E. C. Robinson, “Geometric Deep Learning of the Human Connectome Project Multimodal Cortical Parcellation,” in Machine Learning in Clinical Neuroimaging, 2021, pp. 103–112, doi: 10.1007/978-3-030-87586-2_11.

[33] C. Shain, “CDRNN: Discovering complex dynamics in human language processing,” ACL-IJCNLP 2021 - 59th Annu. Meet. Assoc. Comput. Linguist. 11th Int. Jt. Conf. Nat. Lang. Process. Proc. Conf., pp. 3718–3734, 2021, doi: 10.18653/v1/2021.acl-long.288.

[34] S. M. Kia and A. F. Marquand, “Neural Processes Mixed-Effect Models for Deep Normative Modeling of Clinical Neuroimaging Data,” Proc. Mach. Learn. Res., pp. 297–314, 2018, [Online]. Available: http://arxiv.org/abs/1812.04998.

[35] A. C. Charitos, “Brain disease classification using multi-channel 3D convolutional neural networks,” Linköping University.

[36] S. Yadav, “Bayesian Deep Learning Based Convolutional Neural Network for Classification of Parkinson’s Disease Using Functional Magnetic Resonance Images.”

[37] S. Vieira, W. H. L. Pinaya, and A. Mechelli, “Using deep learning to investigate the neuroimaging correlates of psychiatric and neurological disorders: Methods and applications,” Neuroscience and Biobehavioral Reviews, vol. 74. Elsevier Ltd, pp. 58–75, Mar. 01, 2017, doi: 10.1016/j.neubiorev.2017.01.002.

[38] M. Nazar, M. M. Alam, E. Yafi, and M. M. Su’Ud, “A Systematic Review of Human-Computer Interaction and Explainable Artificial Intelligence in Healthcare with Artificial Intelligence Techniques,” IEEE Access, vol. 9, pp. 153316–153348, 2021, doi: 10.1109/ACCESS.2021.3127881.

[39] J. Gerlings, M. S. Jensen, and A. Shollo, “Explainable AI, But Explainable to Whom? An Exploratory Case Study of xAI in Healthcare,” Intell. Syst. Ref. Libr., vol. 212, no. Ml, pp. 169–198, 2022, doi: 10.1007/978-3-030-83620-7_7.

[40] A. W. Thomas, H. R. Heekeren, K.-R. Müller, and W. Samek, “Analyzing Neuroimaging Data Through Recurrent Deep Learning Models,” Front. Neurosci., Oct. 2019, [Online]. Available: http://arxiv.org/abs/1810.09945.

[41] K. Qiao et al., “Accurate Reconstruction of Image Stimuli From Human Functional Magnetic Resonance Imaging Based on the Decoding Model With Capsule Network Architecture,” Front. Neuroinform., vol. 12, Sep. 2018, doi: 10.3389/fninf.2018.00062.

[42] Y. Yan, E. Solarz, J. Zhu, C. Sripada, M. Duda, and D. Koutra, “Groupinn: Grouping-based interpretable neural network for classification of limited, noisy brain data,” Proc. ACM SIGKDD Int. Conf. Knowl. Discov. Data Min., no. July, pp. 772–782, 2019, doi: 10.1145/3292500.3330921.

[43] X. Li et al., “BrainGNN: Interpretable Brain Graph Neural Network for fMRI Analysis,” Med. Image Anal., vol. 74, p. 102233, 2021, doi: 10.1016/j.media.2021.102233.

[44] Z. Jiang et al., “Attention module improves both performance and interpretability of 4D fMRI decoding neural network,” arXiv, no. Dl.

[45] Y. Du et al., “NeuroMark: An automated and adaptive ICA based pipeline to identify reproducible fMRI markers of brain disorders,” NeuroImage Clin., vol. 28, no. August, p. 102375, 2020, doi: 10.1016/j.nicl.2020.102375.

[46] D. P. Kingma and J. Ba, “Adam: A method for stochastic optimization,” arXiv Prepr. arXiv1412.6980, 2014.

[47] Q. H. Lin, Y. W. Niu, J. Sui, W. Da Zhao, C. Zhuo, and V. D. Calhoun, “SSPNet: An interpretable 3D-CNN for classification of schizophrenia using phase maps of resting-state complex-valued fMRI data,” Med. Image Anal., vol. 79, p. 102430, 2022, doi: 10.1016/j.media.2022.102430.

[48] A. Vilamala, K. H. Madsen, and L. K. Hansen, “Deep convolutional neural networks for interpretable analysis of EEG sleep stage scoring,” IEEE Int. Work. Mach. Learn. Signal Process. MLSP, vol. 2017-Septe, no. 659860, pp. 1–6, 2017, doi: 10.1109/MLSP.2017.8168133.

[49] T. Frick, S. Glüge, A. Rahimi, L. Benini, and T. Brunschwiler, “Explainable Deep Learning for Medical Time Series Data,” in Lecture Notes of the Institute for Computer Sciences, Social-Informatics and Telecommunications Engineering, LNICST, vol. 362 LNICST, 2021, pp. 244–256.

[50] S. Bach, A. Binder, G. Montavon, F. Klauschen, K. R. Müller, and W. Samek, “On pixel-wise explanations for non-linear classifier decisions by layer-wise relevance propagation,” PLoS One,vol. 10, no. 7, Jul. 2015, doi: 10.1371/journal.pone.0130140.

[51] M. Ancona, E. Ceolini, C. Öztireli, and M. Gross, “Towards Better Understanding of Gradient-based Attribution Methods for Deep Neural Networks,” in International Conference on Learning Representations, 2018, pp. 1–16.

[52] W. Samek, A. Binder, G. Montavon, S. Lapuschkin, and K. R. Müller, “Evaluating the visualization of what a deep neural network has learned,” IEEE Trans. Neural Networks Learn. Syst., vol. 28, no. 11, pp. 2660–2673, Nov. 2017, doi: 10.1109/TNNLS.2016.2599820.

[53] W. Yan et al., “Discriminating Schizophrenia From Normal Controls Using Resting State Functional Network Connectivity: A Deep Neural Network and Layer-wise Relevance Propagation Method,” 2017.

[54] C. A. Ellis, R. L. Miller, and V. D. Calhoun, “A Systematic Approach for Explaining Time and Frequency Features Extracted by CNNs from Raw EEG Data,” bioRxiv, 2022.

[55] C. A. Ellis et al., “Novel Methods for Elucidating Modality Importance in Multimodal Electrophysiology Classifiers,” bioRxiv, 2022.

[56] C. A. Ellis, M. S. Sendi, J. T. Willie, and B. Mahmoudi, “Hierarchical Neural Network with Layer-wise Relevance Propagation for Interpretable Multiclass Neural State Classification,” in 10th International IEEE/EMBS Conference on Neural Engineering (NER), 2021, pp. 18–21.

[57] A. Lemay et al., “Monte Carlo dropout increases model repeatability,” arXiv, pp. 1–6, 2021, [Online]. Available: http://arxiv.org/abs/2111.06754.

[58] M. S. E. Sendi et al., “Multiple overlapping dynamic patterns of the visual sensory network in schizophrenia,” Schizophr. Res., vol. 228, pp. 103–111, 2021, doi: 10.1016/j.schres.2020.11.055.

[59] M. Yamamoto et al., “Aberrant functional connectivity between the thalamus and visual cortex is related to attentional impairment in schizophrenia,” Psychiatry Res. - Neuroimaging, vol. 278, no. June, pp. 35–41, 2018, doi: 10.1016/j.pscychresns.2018.06.007.

[60] X. Chen et al., “Functional disconnection between the visual cortex and the sensorimotor cortex suggests a potential mechanism for self-disorder in schizophrenia,” Schizophr. Res., vol. 166, no. 1–3, pp. 151–157, 2014, doi: 10.1016/j.schres.2015.06.014.

[61] M. Ghassemi, L. Oakden-Rayner, and A. L. Beam, “The false hope of current approaches to explainable artificial intelligence in health care,” Lancet. Digit. Heal., vol. 3, no. 11, pp. e745–e750, 2021, doi: 10.1016/S2589-7500(21)00208-9.

